# Colistin resistance and resistance determinants are mobile among *Salmonella enterica* isolates from diseased and healthy pigs in Thailand

**DOI:** 10.1101/2023.01.31.526471

**Authors:** Teerarat Prasertsee, Ben Pascoe, Prapas Patchanee

**Affiliations:** Faculty of Veterinary Science, Prince of Songkla University, Songkhla, Thailand; Centre for Genomic Pathogen Surveillance, Big Data Institute, University of Oxford, Old Road Campus, Oxford, United Kingdom; Ineos Oxford Institute of Antimicrobial Research, Department of Biology, University of Oxford, Oxford, United Kingdom; Integrative Research Center for Veterinary Preventive Medicine, Faculty of Veterinary Medicine, Chiang Mai University, Chiang Mai, Thailand

**Keywords:** *Salmonella*, antimicrobial resistance, multi drug resistant, pig farms, colistin

## Abstract

*Salmonella* is an important enteric pathogen that poses a threat to human and livestock animal health, with emerging multidrug resistance (MDR) a major public health issue globally. We investigated the prevalence of *Salmonella* in healthy and diseased pigs from Thai pig farms and determined their phenotypic and genotypic antimicrobial resistance profiles. A total of 150 fecal samples were collected from pigs housed in pens from four separate pig farms in southern Thailand and tested for the presence of *Salmonella*. Confirmed *Salmonella* isolates were tested for their susceptibility to 11 antimicrobials, and PCR used to detect known antimicrobial resistance genes (ARGs). *Salmonella* isolates were cultured from 69% (103/150) of all fecal samples, with higher prevalence in disease pigs (12/15; 80%), compared with healthy pigs (91/135; 67%). Serotype Rissen was the most frequently identified serotype among the *Salmonella* isolates. Resistance to ampicillin (AMP) (97%), sulfonamide-trimethoprim (SXT) (97%), and tetracycline (TET) (94%) were the most common phenotypes observed. The most common ARGs identified were *bla*_*TEM*_ gene (99.%), *tetA* (87%), *sul1* (77%), and *dfrA1* (74%), and more than 95% of the *Salmonella* isolates tested were MDR - based on resistance to three or more antimicrobial classes. The most common antimicrobial resistance pattern exhibited was AMP-TET-SXT (76%), and resistance to colistin (via the *mcr-1* gene) was observed in both healthy and diseased pigs. The clonal groups of PFGE analysis in serotype Typhimurium revealed the genetic relationship among *Salmonella* isolated from healthy and diseased pigs from different pig farms.

## 1. Introduction

Non-typhoidal *Salmonella* (NTS) are a common cause of human gastroenteritis (Achtman et al., 2012;Phongaran et al., 2019). Infection is usually the result of consumption raw or undercooked food of livestock origin (Sévellec et al., 2020). There has been a dramatic increase in pig production in the 20^th^ Century (Rauw et al., 2020), which has contributed to higher infection levels in livestock pigs (Yang et al., 2019). The detection levels of *Salmonella* in European swine livestock production was 28.5% in 2008, increasing to 66% in 2015 (E.F.S.A, 2017). In south-east Asia, the contamination rates of *Salmonella* infection on livestock farms are very high (Holohan et al., 2022), with prevalence of NTS in pig farms from the Mekong delta region greater than 80% (Tu et al., 2015). In Thailand, the most common *Salmonella enterica* serovars identified from pigs are Rissen and Typhimurium (Phongaran et al., 2019). These serovars commonly infect and cause gastroenteritis in warm blooded animals (Tanner and Kingsley, 2018). Typically, they do not cause severe illness in pigs, symptomatic and asymptomatic animals act as a vehicle, spreading the pathogen to other animals and environments.

Foodborne infection in humans results in gastroenteritis and is usually self-limiting (Holohan et al., 2022). However, antimicrobial treatment is recommended for infections in children under 5 years of age, elderly people, and immunocompromised patients (Sheen et al., 2022). The World Health Organization (WHO) recommended first-choice antibiotic for treatment is ampicillin, trimethoprim-sulfamethoxazole, and chloramphenicol (Tack et al., 2020).

Since 1950, antibiotics such as penicillin, tetracycline, and macrolide have been used in livestock production for increased yield worldwide. Since then, antimicrobial resistance rates have dramatically increased (Holohan et al., 2022). Phenotypic antimicrobial resistance have been linked with veterinary and agricultural use (E.F.S.A et al., 2018) and resistance to one or more antimicrobial agents is commonly found in high rate (74%) (Nüesch-Inderbinen et al., 2019) in isolates from food production animals, especially those form the *Enterobacteriaceae* family (O’Neal et al., 2020). Multi-drug resistance (MDR) in *Salmonella* spp. has been reported in many countries and across the livestock sector, which facilitates the transmission of genes (via horizontal gene transfer) that contribute to acquisition of MDR (Tu et al., 2015; Delgado-Suárez et al., 2019; O’Neal et al., 2020).

Colistin, or polymyxin E, is a bactericidal antibiotic which used as a last-resort drug against infections caused by MDR gram-negative bacteria (Patchanee et al., 2021). Colistin resistance can arise from mutations in chromosomal genes or acquisition of transposable genetic elements carrying mobilized colistin resistance (*mcr*) genes – which result in phenotypic resistance (Holohan et al., 2022). In the *Enterobacteriaceae* family, *mcr-1* and *mcr-3* have been found on the bacterial chromosome, while other *mcr* genes have been identified on plasmids (Ling et al., 2017; Zhou et al., 2017).

The objective of this study was to investigate the prevalence of *Salmonella* on healthy and diseased pigs from pig farms in Southern Thailand and characterise their phenotypic antimicrobial resistance profiles and associated carriage of specific antimicrobial determinants. Our work contributes to surveillance effort in the region and help inform public health interventions for improved food security.

## 2. Materials and Methods

### 2.1. Sample collection and *Salmonella* spp. isolation

A total of 150 samples collected from four pig farms (A-D) in Phatthalung and Songkhla province during 2019 to 2020. The sample population comprised of two groups of pigs: healthy and diseased pigs. Diseased pigs were defined as pigs displaying symptoms of gastrointestinal infection (e.g., watery yellow diarrhea with/without blood or mucus, loss of appetite and fever) and were separated into quarantine stables. The samples were collected from pig feces on the floor in the stable of each group. PSU Animal Standard did not require the study to be reviewed or approved by an Institutional Animal Care and Use Committee because the fecal samples were collected off the ground which non-invasive sampling procedure for animals. All samples were placed in individual sterile packages and transported in insulated boxes with ice to the laboratory within four hours for *Salmonella* identification.

*Salmonella* was isolated using ISO 6579-1:2017 as the standard protocol. Twenty-five-gram samples of fresh fecal samples were placed in sterile bags using aseptic techniques. To each of the samples was added 225 ml of buffered peptone water (BPW; Oxoid, England) and the mixtures were homogenized in a stomacher for 2 min, then incubated for 18 ± 2 h at 37 ± 1°C. Aliquot 3 drops (33 μl each) of incubated BPW were inoculated into a modified semisolid Rappaport Vassiliadis (MSRV; Oxoid, England) medium with novobiocin supplement, and incubated for 24 ± 3 h at 41.5 ± 1°C (negative samples were re-incubated for an additional 24± 3 h). Transferred 1 μl sterile loop of the presumptive *Salmonella*, white/grey turbid zones radiating from the point of inoculation on MSRV, streak onto xylosine lysine deoxycholate (XLD; Oxoid, England) and on brilliant green phenol red lactose sucrose (BPLS; Oxoid, England) incubated for 24 ± 3 h at 37 ± 1°C. Suspected colonies, a red color with a black center on XLD and red and surrounded by a bright red zone on BPLS, were selected to biochemical testing for *Salmonella* spp. confirmation. The serotyping of *Salmonella* was identified at the WHO National *Salmonella* and *Shigella* Center (Nonthaburi, Thailand) following by Kauffmann-White scheme.

### 2.2. Antimicrobial susceptibility testing and minimum inhibitory concentrations

Susceptibility to different antimicrobial agents of all *Salmonella* isolates were performed by disc diffusion method. A total of ten antimicrobials as follows: ampicillin (AMP, 10 μg), amoxycillin/clavulanate (AMC, 20/10 μg), chloramphenicol (CHL, 30 μg), ciprofloxacin (CIP, 5 μg), nalidixic acid (NAL, 30 μg), norfloxacin (NOR, 10 μg), streptomycin (STR 10 μg), sulfisoxazole (SX, 250 μg), tetracycline (TET, 30 μg), and trimethoprim/sulfamethoxazole (SXT, 1.25/23.75 μg) were performed using Mueller-Hinton agar (Oxoid, England). Inhibition zones were measured according to the guidelines of the Clinical and Laboratory Standards Institute (CLSI, 2016). *Escherichia coli* ATCC 25922 was used as an internal quality control.

Gradient diffusion method was used to determine resistance to colistin (COL) according to European Committee on Antimicrobial Susceptibility Testing (EUCAST, 2016) guidelines. Interpretation of colistin resistance were compared to the Annex 1: EUCAST clinical breakpoints and epidemiological cut-off values of Salmonella spp. (MIC dilution ≤ 2 mg/L was recorded as sensitive, while MICs > 2 mg/L were considered resistant).

### 2.3. DNA extraction

All *Salmonella* isolates were grown on nutrient broth (Oxoid, England) at 37 ± 1°C for 24 ± 3 h. Transferred 1 ml of sample to a 1.5 ml microcentrifuge tube for DNA extraction by boiling method. The suspension was centrifuged for 10 min at 14,000×g and discarded the supernatant. The pellet was resuspended in 300 μl of DNase-RNase-free distilled water and mixed by vortex. The tube was centrifuged for 5 min at 14,000×g, and the supernatant was discarded carefully. The pellet was resuspended in 200 μl of DNase-RNase-free distilled water by vortexing. Then, boiling at 100°C for 15 min and immediately chilled on ice. The suspension was centrifuged for 5 min at 14,000×g at 4°C. Transferred the supernatant to a new microcentrifuge tube and boiling at 100°C for 10 min and chilled immediately on ice. The supernatant was used as the DNA template for PCR (Xu et al., 2020).

### 2.4. Detection of antimicrobial resistance genes

All *Salmonella* DNA samples were screened for the presence of antimicrobial resistance genes by PCR using gene-specific primer in table1. PCR reaction was performed at a final volume of 25 ml containing 2 μl of DNA sample, 12.5 μl of 2X Taq Master Mix (Vivantis, Malaysia) which consist of Taq DNA Polymerase (0.05 U/μl), 2X (ViBuffer A, 0.4 mM dNTPs and 3.0 mM MgCl_2_), 1 μl of each primer and 8.5 μl of nuclease-free water. The PCR protocol condition was with initial denaturation step at 94°C for 4 min, followed by 30 cycles at 94°C for 30 sec, 30 sec at primer’s annealing temperature (table1) and 30 sec at 72°C. A final extension step was performed at 72°C for 7 min. and kept on hold at 4°C. Analysis of PCR products were conducted by electrophoresis on 1.0% agarose gel (Agarose Vivantis, Malaysia). DNA bands were visualized by staining with ethidium bromide and photographed under UV illumination.

**Table 1.**
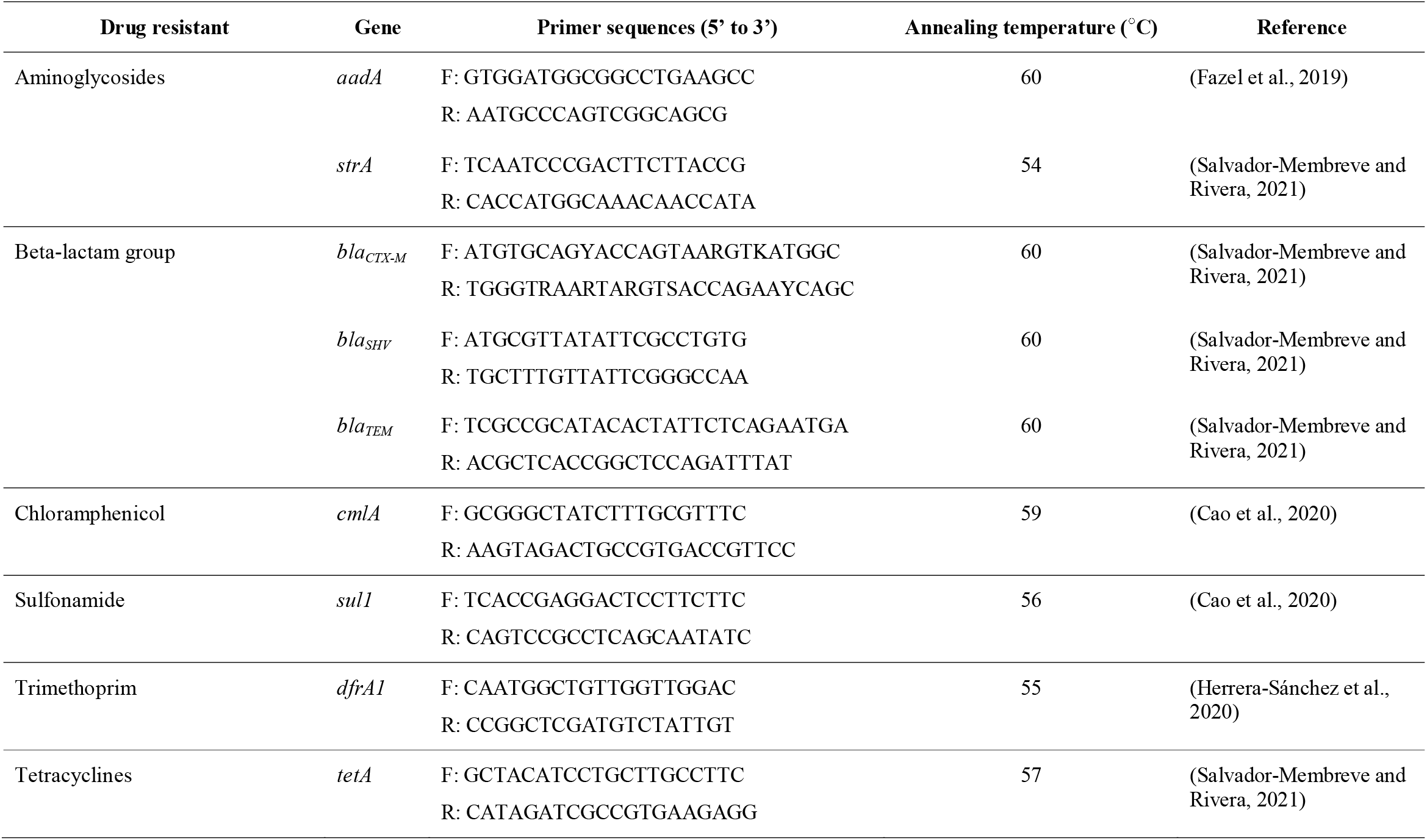
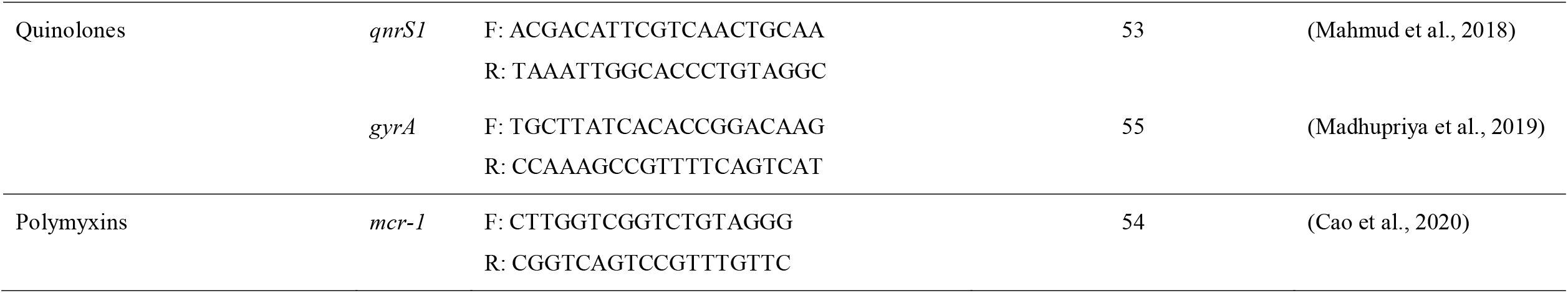
Primers used for detection of genes encoding resistance to different antimicrobials

### 2.5. Pulsed-Field Gel Electrophoresis (PFGE)

Ten *Salmonella* samples resistant to colistin were selected to perform PFGE according to the PulseNet protocol from the Centers for Disease Control and Prevention procedures. In brief, a single colony of each isolate was streaked onto Nutrient agar (Oxoid, England) at 37°C overnight. *Salmonella* colonies were suspended in Cell Suspension Buffer (CBS; 100 mM Tris:100 mM EDTA, pH 8.0) at concentration of optical density (DO) 600. Lysis of bacterial cells used Cell Lysis Buffer (CLB; 50 mM Tris:50 mM EDTA, pH 8.0; 1% Sarcosyl) and Proteinase K (Thermo Fisher Scientific, Korea). Restriction enzyme *Xba*I (Thermo Fisher Scientific, Korea) was used for digestion of genomic DNA samples. The standard strain *Salmonella* Braenderup H9812 (ATCC^®^ BAA-664^™^) was chosen as the universal size standard. Staining gel after electrophoresis used ethidium bromide and sterile ultrapure water was used to destain (Kang et al., 2021).

### 2.6. Statistical and clonal analysis

Association between antimicrobial resistance characteristics and their related resistance genes were analysed by Pearson *Chi*-square of Fisher’s exact tests (where appropriated) in Epi Info™ statistical software using StatCalc modules. The results were considered as statistically significant when p-value < 0.05.

The clonal analysis of ten *Salmonella* DNA fingerprint were analysed by using BioNumerics software version 7.6.3 (Applied Maths, Belgium). The relatedness dendrogram was constructed using Unweighted Pair Group Method with Arithmetic Mean (UPGMA) with Dice coefficient at an optimization setting of 1% and a position tolerance setting of 1%. Interpretation of pulsotype based on the percentage of similarity.

## 3. Results

### 3.1. Prevalence of *Salmonella* positive and serotype distribution

Table 2 presented the prevalence of *Salmonella* positive from pig farms in Patthalung and Songkhla provinces. The overall prevalence was 68.67% (103/150). The highest prevalence of *Salmonella* positive pigs was found on farm B (84.00%) followed by farm C (78.00%), farm D (64.00%), and farm A (54.00%). *Salmonella* was just as likely to be sampled from healthy pigs, with successful identification of isolates from 82.61%, 80.00%, 60.00% and 51.06% of pigs from farms B, C, D and A, respectively. *Salmonella* isolates were identified from all diseased pigs (100%) from farms A and B, and 80% and 60% of pigs from farms D and farm C, respectively.

**Table 2.** Prevalence of *Salmonella* positive from pig farms in Phatthalung and Songkhla provinces

| Farm |  | No. of<br>samples | No. of<br>positive<br>samples | Prevalence of positive samples<br>(95% CI) |
| --- | --- | --- | --- | --- |
| A<br>(Songkhla) | Healthy pig | 47 | 24 | 51.06 (36.77-65.36) |
|  | Diseased pigs | 3 | 3 | 100.00 (100.00-100.00) |
|  | **Total** | **50** | **27** | **54.00 (40.19-67.81)** |
| B<br>(Phatthalung) | Healthy pig | 23 | 19 | 82.61 (67.12-98.10) |
|  | Diseased pigs | 2 | 2 | 100.00 (100.00-100.00) |
|  | **Total** | **25** | **21** | **84.00 (69.63-98.37)** |
| C<br>(Phatthalung) | Healthy pig | 45 | 36 | 80.00 (68.31-91.69) |
|  | Diseased pigs | 5 | 3 | 60.00 (22.91-88.40) |
|  | **Total** | **50** | **39** | **78.00 (66.52-89.48)** |
| D<br>(Songkhla) | Healthy pig | 20 | 12 | 60.00 (38.60-78.17) |
|  | Diseased pigs | 5 | 4 | 80.00 (35.96-97.97) |
|  | **Total** | **25** | **16** | **64.00 (45.18-82.82)** |
| **Total** |  | **150** | **103** | **68.67 (61.24-76.09)** |

The serotype distribution of *Salmonella* isolates is shown in table 3, with eighteen different serotypes observed in this study. Rissen (n=21) was the most common serotype found within pig farms, followed by serotypes Typhimurium (n=17), Enteritidis (n=11) and Stanley (n=10). Fourteen serotypes were observed in only healthy pigs while four serotypes; *S*. Typhimurium, *S*. Enteritidis, *S*. Brunei, and *S*. 4,5,12:i:-, were isolated from both healthy and diseased pigs. The dominant serotypes from healthy and diseased pigs were *S*. Rissen (n=21) and *S*. 4,5,12:i:- (n=5), respectively.

**Table 3.**
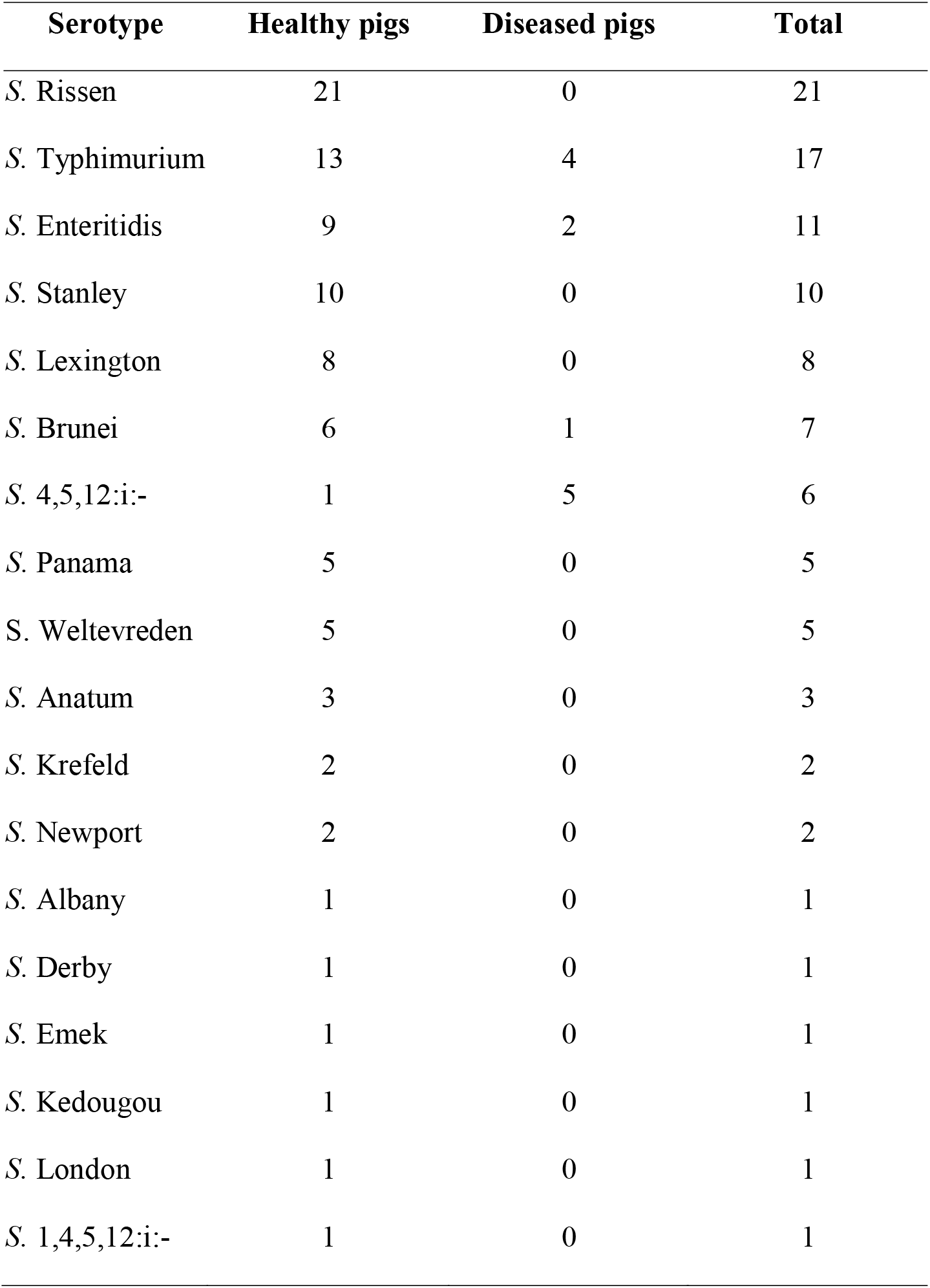
Serotypes distribution of *Salmonella* isolated from healthy and diseased pigs.

### 3.2. Antimicrobial resistance of *Salmonella* isolates

Table 4 showed the results of antimicrobial resistance genes among the 103 *Salmonella* isolates. The presence of twelve known resistance genes were assessed, from eight different categories of antimicrobials including aminoglycosides (*aadA, strA*), beta-lactams (*bla*_*CTX-M*_, *bla*_*SHV*_, *bla*_*TEM*_), chloramphenicol (*cmlA*), sulfonamide (*sul1*), trimethoprim (*dfrA1*), tetracyclines (*tetA*), quinolones (*qnrS1, gyrA*), and polymyxins (*mcr-1*). The beta-lactam resistance gene, *bla*_*TEM*,_ was the most prevalent resistance gene, found in 99.03% (102/103) of isolates - both healthy and diseased pigs (table 4). The *bla*_*SHV*_ gene, from the same group, was not found in any of the isolates. High rates of prevalence were observed for the antimicrobial resistance genes: *tetA* (87.38%; 90/103), *sul1* (76.70%; 79/103), and *dfrA1* (73.79%; 76/103), and were observed in both healthy and diseased pigs (table 4). Carriage of the *mcr-1* gene was identified in 15.53% (16/103) of isolates, of which most were healthy pigs (n=9 out of 16).

**Table 4.** Antimicrobial resistance genes of *Salmonella* isolates from healthy and diseased pigs

| Resistance genes | No. of positive resistance gene (%) |  |
| --- | --- | --- |
|  | Healthy pigs | Diseased pigs |
| *aadA* | 3 (2.91) | 1 (0.97) |
| *strA* | 8 (7.77) | 3 (2.91) |
| *bla<sub>CTX-M</sub>* | 2 (1.94) | 0 (0.00) |
| *Bla<sub>SHV</sub>* | 0 (0.00) | 0 (0.00) |
| *bla<sub>TEM</sub>* | 90 (87.38) | 12 (11.65) |
| *cmlA* | 5 (4.85) | 0 (0.00) |
| *sulI* | 69 (66.99) | 10 (9.71) |
| *dfrA1* | 67 (65.05) | 9 (8.74) |
| *tetA* | 79 (76.70) | 11 (10.68) |
| *qnrS1* | 3 (2.91) | 3 (2.91) |
| *gyrA* | 4 (2.88) | 1 (0.97) |
| *mcr-1* | 9 (8.74) | 7 (6.80) |

Summary results of antimicrobial susceptibility testing among *Salmonella* isolates are shown in table 5. All 103 *Salmonella* isolates were susceptible to AMC, CHL and CIP. High rates of resistance were observed for AMP and SXT (97.09%; 100/103), followed by TET (94.17%; 97/103), COL (9.71%; 10/103), STR (7.77%; 8/103), SX (5.83%; 6/103), NAL (3.88%; 4/103), and NOR (1.94%: 2/103). 88.44% of healthy pig isolates was resistant to AMP and SXT while 11.65% of diseased pig isolates was resistant to AMP, TET and SXT. Polymyxins resistance was low in both healthy pigs (3.88%) and diseased pigs (5.83%) (table 5).

**Table 5.** Antimicrobial resistance genes of *Salmonella* isolates from healthy and diseased pigs

| Antimicrobial agents | No. of resistance (%) |  |
| --- | --- | --- |
|  | Healthy pigs | Diseased pigs |
| AMP | 88 (88.44) | 12 (11.65) |
| AMC | 0 (0.00) | 0 (0.00) |
| CHL | 0 (0.00) | 0 (0.00) |
| CIP | 0 (0.00) | 0 (0.00) |
| NAL | 2 (1.94) | 2 (1.94) |
| NOR | 1 (0.97) | 1 (0.97) |
| STR | 5 (4.85) | 3 (2.91) |
| SX | 4 (3.88) | 2 (1.94) |
| TET | 85 (82.52) | 12 (11.65) |
| SXT | 88 (85.44) | 12 (11.65) |
| COL | 4 (3.88) | 6 (5.83) |

The antimicrobial resistance patterns of all *Salmonella* isolates were presented in table 6, and the *S*. 1,4,5,12:i:- (ID: SK_D_PH_SL4) isolated from healthy pigs in farm D in Songkhla province was sensitive to all eleven antimicrobial agents tested. One *S*. Typhimurium isolate from healthy pigs (ID: PL_B_PH_N3) was resistant to a single antimicrobial (SXT). A small number of isolates from healthy pigs were resistant to two antimicrobial classes, demonstrating and the antimicrobial resistance patterns AMP-TET (n=1) and AMP-SXT (n=2). Multidrug resistance (MDR) was defined as resistance to at least one agent from three or more different antimicrobial classes, and thirteen different MDR patterns of resistance were observed (table 6). In total, 95.15% (98/103) of *Salmonella* isolates were found to be MDR. Eighty-six from ninety-one of *Salmonella* isolates from healthy pigs were MDR. All twelve *Salmonella* isolates from diseased pigs were identified as MDR, and half of them demonstrated the same MDR pattern (AMP-TET-SXT) (table 6). The highest prevalence MDR pattern was AMP-TET-SXT (75.73%; 78/103), followed by AMP-TET-SXT-COL (3.88%; 4/103), and AMP-STR-TET-SXT (2.91%; 3/103). Three MDR patterns including AMP-TET-SXT, AMP-TET-SXT-COL, and AMP-SX-TET-SXT-COL were demonstrated in *Salmonella* isolates sampled from both healthy and diseased pigs. Additionally, The MDR patterns presented only in *Salmonella* isolates from diseased pigs comprised of AMP-NAL-TET-SXT-COL (n=1), AMP-STR-TET-SXT-COL (n=1), AMP-STR-SX-TET-SXT-COL (n=1), and AMP-NAL-NOR-STR-TET-SXT-COL (n=1). However, resistance to six different antimicrobial classes and seven antimicrobial agents (AMP-NAL-NOR-STR-TET-SXT-COL) was observed in *S*. 4,5,12:i:-from diseased pigs (ID: PL_C_PD_1).

**Table 6.** Antimicrobial resistance pattern of *Salmonella* isolates from healthy and diseased pigs

| Antimicrobial resistance patterns | No. of isolates (%) |  |
| --- | --- | --- |
|  | Healthy pigs | Diseased pigs |
| **Pan-susceptible** | 1 (0.97) | 0 (0.00) |
| **Single-drug resistance** |  |  |
| SXT | 1 (0.97) | 0 (0.00) |
| **Two-drug resistance** |  |  |
| AMP-TET | 1 (0.97) | 0 (0.00) |
| AMP-SXT | 2 (1.94) | 0 (0.00) |
| **Multidrug resistance** |  |  |
| NOR-SX-SXT | 1 (0.97) | 0 (0.00) |
| AMP-STR-TET | 1 (0.97) | 0 (0.00) |
| AMP-STR-SXT | 1 (0.97) | 0 (0.00) |
| AMP-TET-SXT | 72 (69.90) | 6 (5.83) |
| AMP-SX-TET-SXT | 2 (1.94) | 0 (0.00) |
| AMP-NAL-TET-SXT | 2 (1.94) | 0 (0.00) |
| AMP-STR-TET-SXT | 3 (2.91) | 0 (0.00) |
| AMP-TET-SXT-COL | 3 (2.91) | 1 (0.97) |
| AMP-SX-TET-SXT-COL | 1 (0.97) | 1 (0.97) |
| AMP-NAL-TET-SXT-COL | 0 (0.00) | 1 (0.97) |
| AMP-STR-TET-SXT-COL | 0 (0.00) | 1 (0.97) |
| AMP-STR-SX-TET-SXT-COL | 0 (0.00) | 1 (0.97) |
| AMP-NAL-NOR-STR-TET-SXT-COL | 0 (0.00) | 1 (0.97) |

### 3.3. Association between antimicrobial resistance phenotype and corresponding genes

Statistical analysis of the antimicrobial resistance phenotypes and the corresponding genes revealed that AMP and *bla*_*TEM*_ gene, SXT and *sul1* gene, TET and *tetA* gene, COL and *mcr-1* gene, STR and *aadA* gene, STR and *strA* gene, NAL and *qnrS1* gene, NAL and *gyrA* gene, NOR and *qnrS1* gene, and NOR and *gyrA* gene were significantly related to the observed resistance (p-value<0.05) (figure 1).

**Figure 1.**
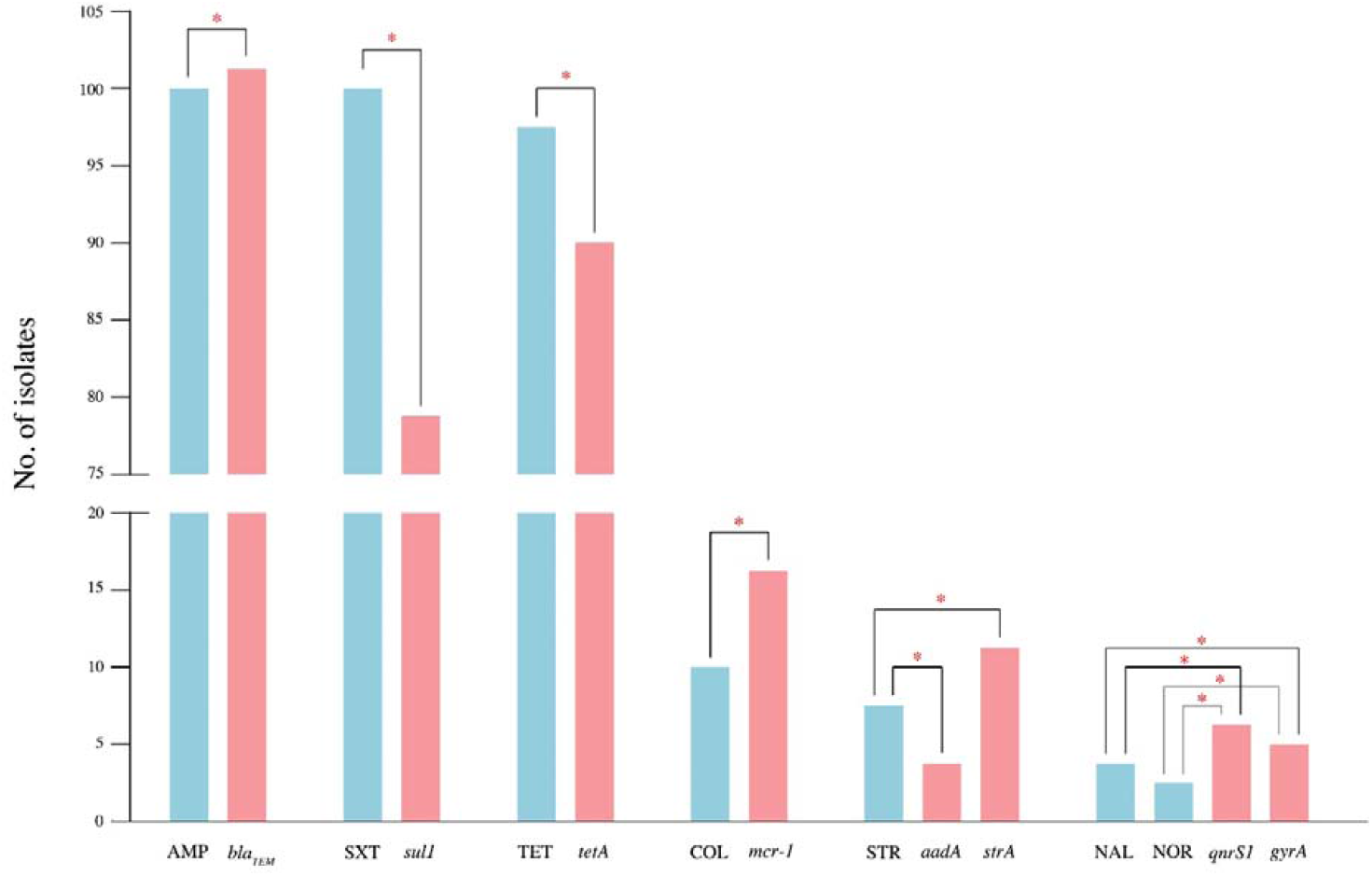
The relationship between antimicrobial resistance phenotype (blue bars) and related genes (pink bars) of all 103 *Salmonella* isolates. Ten resistance phenotype and twelve resistance genes (AMP and *bla*_*TEM*_, *bla*_*CTX-M*_, *bla*_*SHV*_; AMC and *bla*_*TEM*_, *bla*_*CTX-M*_, *bla*_*SHV*_; CHL and *cmlA;* CIP and *qnrS1, gyrA;* NAL and *qnrS1, gyrA;* NOR and and *qnrS1, gyrA;* STR and *aadA, strA;* SX and *sul1;* SXT and *sul1, dfrA1;* TET and *tetA;* COL and *mcr-1*) were analysed. The significantly relatedness (p-value<0.05) between antimicrobial resistance characteristics and their genes presented in this graph.

### 3.4. Clonal relatedness

All ten *Salmonella* isolates that demonstrated colistin resistance were selected for PFGE, including isolates from *S*. Typhimurium (n=4), *S*. 4,5,12:i:- (n=3), *S*. Rissen (n=2), and *S*. Weltevreden (n=1) serovars. Within each serovar, isolates grouped into clonal clusters of pulsotypes (figure 2). Cluster 1 belonged to *S*. 4,5,12:i:-isolated from diseased pigs in farm C and D. The antimicrobial resistance profile of three isolates was different within cluster 1. Four *S*. Typhimurium were grouped into cluster 2. The isolates in this cluster were from both healthy pig and diseased pigs and carried the same resistance genes profile (figure 2). The identical clone of two *S*. Rissen from healthy pigs was in cluster 3. Although *S*. Rissen in this group isolated from different farms (B and D), these two isolates had the same antimicrobial resistance characteristics (figure 2). *S*. Weltevreden from healthy pig in farm D was classified into cluster 4.

**Figure 2.**
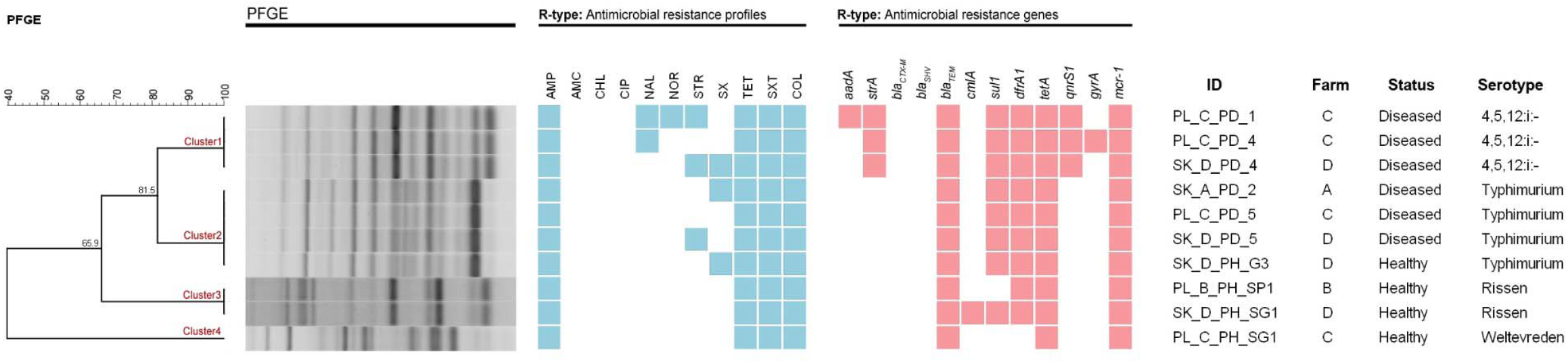
Phylogenetic tree of DNA fingerprint of ten *Salmonella* isolates with R-type of antimicrobial resistance phenotype and genotype. Pink box = resistance phenotype, blue box = resistance genotype.

## 4. Discussion

In this study, the prevalence of *Salmonella* in farmed pigs was 68.67% (95%CI: 61.24-76.09), which is higher than many recent the reports (Casanova-Higes et al., 2019), including those from Thailand. Recent Thai estimated reported *Salmonella* prevalence of 6% (Padungtod and Kaneene, 2006), 34.98% (Tadee et al., 2014) and 21.51% (Anuchatkitcharoen et al., 2020). The differences in prevalence could be a result of differences in farm management strategies, sanitation, hygiene practices, and the individual environmental conditions of each farm (Cao et al., 2020). Furthermore, fifteen isolates in this study collected from feces of pigs that had gastrointestinal problems such as diarrhea in quarantine housing. Specific sampling from diseased pigs may have contributed to the higher prevalence observed in this study.

In this study, Rissen was the dominant serotype on the pig farms, consistent with several other studies reporting serotype Rissen as the most common serotype found in pig production in Thailand (Patchanee et al., 2016;Yang et al., 2018;Phongaran et al., 2019). Despite this, there have been few reports of this serotype from pig farms in Southern Thailand. In 2013, Rissen was found as the predominant serotype isolated from pork in Phatthalung province (Southern Thailand) (Lertworapreecha et al., 2013). Taken together, our findings suggest that Rissen is a common serovar infecting pigs from the region. Although this serovar is thought to have emerged in SE Asia, it has likely spread around the world though global pork markets (Elbediwi et al., 2021) and the European Food Safety Authority (EFSA) and European Centre for Disease Prevention and Control (ECDC) reported it among the top twenty *Salmonella* serovars linked with human salmonellosis (E.F.S.A, 2017; Xu et al., 2020).

Although NTS rarely cause severe illness in healthy pigs, presenting as gastroenteritis and diarrhea, which effects pig growth. Consequently, antimicrobials are often used prophylactically to prevent disease in asymptomatic animals, with ampicillin and tetracycline the preferred antimicrobials to treat diarrheal symptoms (Rosager et al., 2017). This was reflected in our results, with extensive resistance to AMP (97.09%), SXT (97.09%), and TET (94.17%) reported. Previous studies in the north-east of Thailand and Laos have also observed extremely high levels of resistance to AMP (100.00%), SXT (100.00%), and TET (91.40%) (Sinwat et al., 2016). This may be attributed to the common usage of the antimicrobial agents in livestock production.

Resistance to more than one antimicrobial agent was observed in 98.06% of *Salmonella* isolates, and approximately 95% were MDR. This is consistent with the current crisis in south-east Asia. MDR *Salmonella* have been isolated from livestock production animals in extremely high numbers, including 98% in Thailand, 98.4% Laos, 59.4% in Vietnam and 52% in Cambodia (Tu et al., 2015; Sinwat et al., 2016; Trongjit et al., 2017; Holohan et al., 2022). The ubiquitous prevalence of MDR isolates from pigs in this region is alarming and poses a current public health risk.

In the livestock farming sector, especially pig production, antibiotics have been used extensively for treatment, prophylaxis, metaphylaxis and growth promotion. This pervasive use of antibiotics has contributed to a genetic bottleneck that promotes the spread and dissemination of resistant genotypes among enteric bacteria such *as Salmonella* (Cao et al., 2020). However, several resistance phenotypes require multiple genes, as highlighted in our study where several genetic determinants showed little association with their predicted resistance phenotype (figure 1). In this study, the most common antimicrobial resistant gene was *bla*_*TEM*_ gene (99.03%) and correlated well with ampicillin (AMP) resistance. An additional two isolates carried the *bla*_*CTX-M*_ gene, but no isolate harboured the *bla*_*SHV*_ gene. The *bla*_*CTX-M*_ and *bla*_*SHV*_ genes are typically the most common extended-spectrum β-lactamase (ESBL) genes, and strains carrying these genes often demonstrate resistance to additional antimicrobial agents belonging to other classes (Pishtiwan and Khadija, 2019). In our collection, two *S*. Typhimurium isolates harboured the *bla*_*CTX-M*_ gene and were also found to be resistant to multiple antibiotic classes. Tetracycline resistance genes (*tetA*) and sulfonamide-trimethoprim (*sul1* and *dfrA1*) were also detected in high frequency. These genes were also common in other studies from the region (Lapierre et al., 2008; Thai et al., 2012; Prasertsee et al., 2019; Karabasanavar et al., 2022), reflected in the high rates of resistance to β-lactam, tetracycline, and sulfonamide-trimethoprim.

Colistin resistance in *Enterobacteriaceae* family bacteria has been reported globally. The *mcr-1* gene plays an important role in colistin resistance of *Salmonella* isolates from livestock animals, environmental and human (Lu et al., 2018; Cao et al., 2020). In this study, sixteen isolates contained the *mcr-1* gene, ten of which demonstrated correlating phenotypic colistin resistance. Six colistin resistant samples were obtained from symptomatic pigs (diarrhoea) in farm A (n=1), C (n=3) and D (n=2) and other four isolates were collected from healthy pigs in farm B (n=1), C (n=1) and D (n=2). On farm C and D, the symptomatic pigs were treated by mixing the colistin in animal feed and continued feeding until recovery. This was the general practice in the small-scale farms of this area. Use of antimicrobials in animal feed for treatment can expose bacteria to antibiotics at sub-therapeutic levels, which may contribute to increasing the selective pressure on the wider bacterial population.

The DNA fingerprint of ten *Salmonella* isolates resistant to colistin was analysed and *Salmonella* isolates were grouped according to serotype in clonal clusters 1-4 (figure 2). The clonal groups of each serotype indicated that *Salmonella* isolates shared similar genetic backgrounds. Overall, diseased pigs were resistant to a broader spectrum of antibiotics, potentially due to the likely increased exposure to antimicrobial agents. It was therefore interesting to note that healthy pigs also represent a potential reservoir of MDR. This has been noted with other species, with healthy pigs harbouring diverse resistant genotypes of *Streptococcus suis* (Kittiwan et al., 2022).

## 5. Conclusion

In conclusion, this study confirmed that the *Salmonella* can be found and circulated within pig farm base on the evidence of high prevalence rate of *Salmonella* positive from both healthy and diseased pigs. High rates of MDR *Salmonella* were observed, especially in diseased pigs with more than half of the collected isolates resistant to three or more antibiotic classes. Taken with evidence from other studies, this poses a current public health risk in the region. Preventive control strategies must account for these extremely high levels of antibiotic resistance to stop the spread of MDR and reduce public exposure to MDR isolates via the pork production chain.

## 6. Conflict of Interest

The authors declare that the research was conducted without commercial or financial relationships that could be construed as a potential conflict of interest.

## 7. Author Contributions

TP collected samples, conducted microbiology and molecular analysis, and wrote the manuscript. BP and PP helped conceive the study, provided guidance and edited the manuscript.

## 8. Funding

This study was financially supported by Prince of Songkla University (Grant No. VET6302139N) and partially funded by Chiang Mai University. The authors would like to acknowledge funding from the Medical Research Council (UK; MR/V001213/1) and the National Institute of Allergy and Infectious Diseases (1R21AI163801-01A1 & 5R01AI158576-02).

## Notes

### Competing Interest Statement

The authors have declared no competing interest.

